# Second-Generation Antibodies Neutralize Emerging SARS-CoV-2 Variants of Concern

**DOI:** 10.1101/2021.06.09.447527

**Authors:** Branislav Kovacech, Lubica Fialova, Peter Filipcik, Rostislav Skrabana, Monika Zilkova, Natalia Paulenka-Ivanovova, Andrej Kovac, Denisa Palova, Gabriela Paulíkova Rolkova, Katarina Tomkova, Natalia Turic Csokova, Karina Markova, Michaela Skrabanova, Kristina Sinska, Neha Basheer, Petra Majerova, Jozef Hanes, Vojtech Parrak, Michal Prcina, Ondrej Cehlar, Martin Cente, Juraj Piestansky, Michal Fresser, Michal Novak, Monika Slavikova, Kristina Borsova, Viktoria Cabanova, Bronislava Brejova, Tomas Vinař, Jozef Nosek, Boris Klempa, Norbert Zilka, Eva Kontsekova

**Author notes:** Correspondence should be addressed to B.K or N.Z., Branislav Kovacech, PhD, AXON Neuroscience R&D Services SE, Dvorakovo nabrezie 11, 811 02, Bratislava, Slovakia, Norbert Zilka, PhD, AXON Neuroscience R&D Services SE, Dvorakovo nabrezie 11, 811 02, Bratislava, Slovakia.

## Abstract

Recently emerged SARS-CoV-2 variants show resistance to some antibodies that were authorized for emergency use. We employed hybridoma technology combined with authentic virus assays to develop second-generation antibodies, which were specifically selected for their ability to neutralize new variants of SARS-CoV-2. AX290 and AX677, two monoclonal antibodies with non-overlapping epitopes, exhibit subnanomolar or nanomolar affinities to the receptor binding domain of the viral Spike protein carrying amino acid substitutions N501Y, N439K, E484K, K417N, and a combination N501Y/E484K/K417N found in the circulating virus variants. The antibodies showed excellent neutralization of an authentic SARS-CoV-2 virus representing strains circulating in Europe in spring 2020 and also the variants of concern B.1.1.7 and B.1.351. Finally, the combination of the two antibodies prevented the appearance of escape mutations of the authentic SARS-CoV-2 virus. The neutralizing properties were fully reproduced in chimeric mouse-human versions, which may represent a promising tool for COVID-19 therapy.

## Introduction

Neutralizing antibodies (NAbs) against SARS-CoV-2 are being evaluated for prophylaxis and as therapeutic agents for COVID-19 patients^1^. Human antibodies isolated from COVID-19 convalescent patients often target the receptor-binding domain (RBD) of the Spike (S) protein of SARS-CoV-2, and recognize distinct, sometimes non-overlapping epitopes ^2–8^. Only a small subset of these antibodies can block viral entry, usually by interfering with the binding of the viral S protein to the cellular receptor ACE2 ^2,4,5^. Clinical trials suggest that antibody treatments can prevent deaths and hospitalizations among people with mild or moderate COVID‑19. Some of therapeutic antibodies against COVID-19 were already authorized for emergency use ^9^, however they were not initially selected for the neutralization activity on emerging variants of COVID-19.

The majority of neutralizing antibodies against SARS-CoV-2 in the clinical development was derived from convalescent patients who recovered from COVID-19 ^10–13^. Antibodies derived from human B cells do not require an extensive humanization process and can be fast-forwarded to manufacture ^14^. However, several studies demonstrated that neutralizing monoclonal antibodies obtained from individuals during the early convalescence period may exhibit low levels of maturation associated with viral suppression of germinal center development ^15,16^. Near-germline human antibodies could cross react with antigens in mammalian tissues ^17^. Optimal immunization procedure and exploration of mouse immunoglobulin repertoire may overcome these limitations and survey an orthogonal conformation space of antibody specificities ^18^.

The emergence of new SARS-CoV-2 variants of concern (VOCs) B.1.1.7 (South East of England) ^19,20^, B.1.351 (South Africa) ^21^, and P.1 (Brazil) ^22,23^ that harbor mutations in the viral S protein raised alarm whether current vaccines and therapeutic antibodies were sufficiently effective. Indeed, several independent studies have shown that mutant variants are partially or completely resistant against therapeutic antibodies that were authorized for emergency use ^24–28^.

Here we present two mouse-human chimeric monoclonal antibodies against the S protein RBD with non-overlapping epitopes and excellent neutralization potency not only against a live authentic SARS-CoV-2 virus isolate that circulated in Europe in spring 2020, but also variants of concern B.1.1.7 and B.1.351 currently spreading in the population. The combination of the two antibodies prevents generation of escape mutations of the live authentic SARS-CoV-2 virus, making the mixture a promising candidate for a therapeutic application.

## Results

### Identification of monoclonal antibodies neutralizing SARS-CoV-2

To isolate fully matured antibodies with high neutralizing potency against live SARS-CoV-2 virus, we immunized mice either with the recombinant Spike protein or its RBD. Screening of the best antibody-expressing mouse hybridoma clones generated from mice immunized with the viral Spike protein revealed that all except one (98%) target RBD of the SARS-CoV-2 S protein (Figure 1A). Nineteen out of 23 hybridoma clones from RBD-immunized mice produced antibodies recognizing the S protein (Figure 1B). Out of the 65 RBD- and Spike-positive hybridoma clone supernatants, thirteen inhibited the interaction between RBD and ACE2 in the competition ELISA, twelve inhibited the interaction between the S protein and ACE2 expressed on the surface of HEK293T/17-hACE2 cells (the assay mimics the first step in the infection of the human cells by SARS-CoV-2), twelve inhibited cell entry of replication-deficient mouse leukemia virus (MLV) pseudotyped with the SARS-CoV-2 spike protein, and thirteen neutralized the live authentic virus (Slovakia/SK-BMC5/2020 isolate, B.1 lineage) in PRNT (Figure 1C).

**Figure1.**
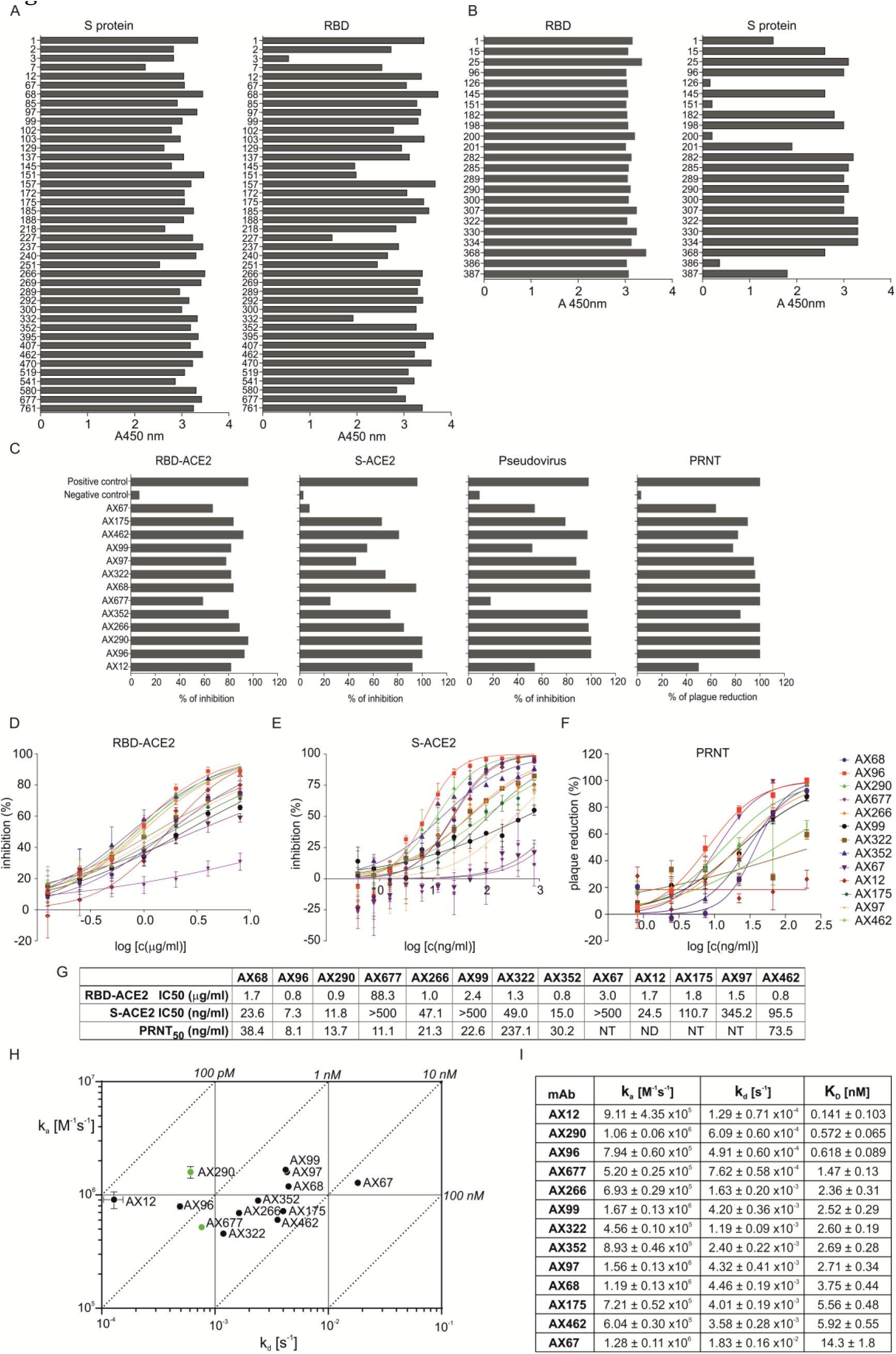
Binding characteristics and inhibition activities of monoclonal antibodies specific to SARS-CoV-2 S/RBD. (A) ELISA immunoreactivity of hybridoma clones derived from mice immunized with the Spike (S) protein of SARS-CoV-2 and with RBD of Spike protein (B). (C) Blocking of interaction between RBD and ACE2 in a competition assay based on ELISA (RBD-ACE2), inhibition of the S protein internalization by cells expressing ACE2 (S-ACE2), inhibition of cell infection by MLV pseudotyped with SARS-CoV-2 spike protein (Pseudovirus) and plaque reduction neutralization test performed with an authentic SARS-CoV-2 virus isolate Slovakia/SK-BMC5/2020 (PRNT). Experiments were done with hybridoma culture supernatants adjusted with fresh cell culture medium to the same mAb concentration and then diluted for the assays as follows: RBD-ACE2 competitive ELISA 1:6 in PBS-T; S-ACE2 cell assay 1:50 in DMEM; pseudoviral assay 1:25 in DMEM; live virus PRNT 1:50 in EMEM. Positive control: serum of a mouse immunized with the S protein, diluted 1:200; negative control: irrelevant mAb. (G-J). All experiments are an average of at least two measurements, pseudoviral tests were all done in tetraplicates. (D) The purified monoclonal antibodies were tested for the inhibition of ACE2-mediated S protein internalization by HEK293/17-hACE2 cells, blocking the RBD-ACE2 interaction in competitive ELISA (E) and plaque reduction neutralization test (F) performed with strain Slovakia/SK-BMC5/2020. The curves are calculated from 2 replicates using Prism 6 for Windows (GraphPad Software). (G) Summary of assays from D-F. (H, I) Kinetic characteristics of the interactions of RBD with selected neutralizing antibodies. On-rate (k_a_), off-rate (k_d_) constants and equilibrium dissociation constant (K_D_) of individual antibody-RBD complexes (±SD). Green dots mark antibodies AX290 and AX677 selected for further development.

Quantitative analyses of purified 13 best-performing monoclonal antibodies (mAbs) revealed that the majority of them inhibited the interaction of the RBD and ACE2 proteins in competitive ELISA with IC50 of 0.8 – 3.0 μg/ml (Figure 1D, G). MAbs AX96, AX290 and AX352 had the strongest inhibitory effect on the binding and internalization of the S protein by HEK293T/17-hACE2 cells (Figure 1E, G), with IC50 values of 7.3 ng/ml, 11.8 ng/ml and 15.0 ng/ml, respectively. Five antibodies displayed high neutralization potency of the live SARS-CoV-2 virus in PRNT (Figure 1F, G). Importantly, AX677 showed the second highest live virus neutralization activity of all tested antibodies (PRNT_50_=11.1 ng/ml), although it had relatively limited ability to inhibit Spike-ACE2 interaction in *in vitro* assays and poorly inhibited the S-typed MLV pseudovirus.

The neutralizing antibodies bind RBD with nanomolar or sub-nanomolar affinities (Figure 1H). Kinetic SPR experiments revealed that antibodies exhibited a large distribution of dissociation kinetic rates, spanning two orders of magnitude (1.3×10^−4^ – 1.8×10^−2^ s^−1^; Figure 1H, I). The association kinetics also varied, but to a lesser extent (4.6×10^5^ – 1.7×10^6^ M^−1^s^−1^). Four antibodies with the highest (picomolar) affinities, AX12, AX96, AX290 and AX677, had very slow dissociation rates below 10^−3^ s^−1^. Six antibodies showed nearly identical affinities of 2 to 3 nM (AX68, AX97, AX99, AX266, AX322 and AX352), but with contrasting kinetic behaviour. In general, the antibodies exhibit RBD binding affinities similar to or better than antibodies isolated from patient sera ^12,24,25^.

### Neutralizing antibodies recognize two immunodominant epitopes on RBD

Mutual competition of the selected antibodies for binding to RBD revealed three non-overlapping antibody epitopes (Figure 2A, Epitopes I, II, III). Epitope I was recognized by the group of six antibodies, which appeared among the best neutralizers of the authentic live SARS-CoV-2 virus. Within this group, the virus neutralization activities highly correlated with their affinities to RBD (R^2^>0.9). The Epitope II group comprised six antibodies, where AX677 and AX175 did not compete with each other. One antibody (AX12) recognised a non-overlapping independent region (Epitope III).

**Figure 2.**
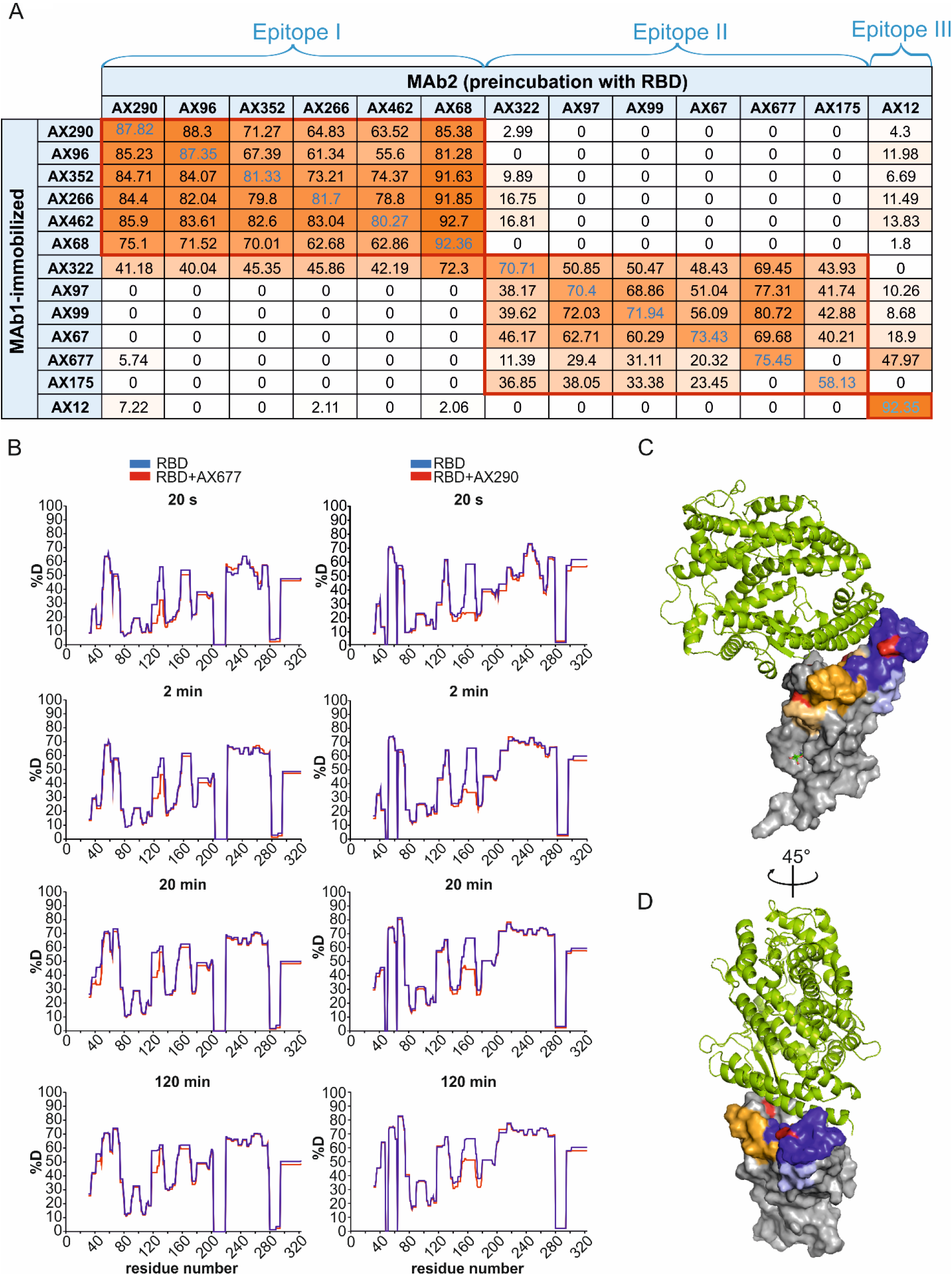
The panel of selected neutralization antibodies target three non-overlapping epitopes on RBD. (A) Competitive ELISA was used for determination of the epitopes on RBD by neutralizing mAbs. Signal reduction for more than 30 % was considered a positive competition. (B) Difference plots depicting the changes in deuterium uptake in the RBD peptides upon binding to AX677 (left panels) and AX290 (right panels). The HDX reaction times are indicated in seconds. Numbers on x-axis represent amino acid positions of the recombinant RBD protein (adding 316 aligns them with the S protein numbering). Each plot represents an average of three technical replicates. (C, D) The positions of the peptides bound by the antibodies and identified in the HDX experiments are highlighted in the structure of SARS-CoV-2 Spike receptor-binding domain bound to ACE2 reproduced from PDB 6M0J (*34*). The AX290 binding site is represent by shades of blue, AX677 by orange and yellow. Darker colours represent peptides that exhibit reduction in deuterium exchange (%D) ≥20%, light colours represent peptides with reduction in %D between 5-20%. ACE2 is shown in a green cartoon model, RBD as a grey surface model, mutations N439K (within AX677 binding site) and E484K (within AX290 binding site) are shown in red. The model was rendered using the PyMOL Molecular Graphics System, Version 2.5.0a0 Open-Source, Schrödinger, LLC.

AX290 and AX677 showed the highest binding affinities to RBD of all members of the Epitope I and II groups, respectively, and showed the highest neutralizing activities against the live authentic virus. We applied hydrogen-deuterium exchange coupled to mass spectrometry (HDX) to identify their binding sites on RBD. Deuterium uptake was monitored on 288 unique peptides, covering 89% of the RBD sequence. The peptides protected by AX677 antibody included amino acids 42-48 and 118-139 (RBD numbering) (Figure 2B), which correspond to amino acids 358-364 and 434-455, respectively, of the S protein. The RBD peptides protected by the AX290 antibody encompassed amino acids 151-179 (Figure 2B), which correspond to peptides 467-495, respectively, of the S protein. We used the published model of ACE2-bound RBD from PDB 6M0J^29^ and highlighted the peptides protected in HDX experiments (Figure 2C, D). The epitope of AX290 is located in the region of RBD that comprises some of the contacts to ACE2, is often targeted by neutralizing Abs ^2^, and overlaps with epitopes of REGN10933 and LY-CoV555^12,30^. The epitope of AX677 is located outside of the ACE2-binding interface, which explains why it does not inhibit ACE2-RBD interaction. Its mode of neutralization might be allosteric or via interference with attachment to other cell surface molecules.

### Selection of antibodies that neutralize SARS-CoV-2 variants of concern B.1.1.7 and B.1.351

In order to test the ability of the selected mAbs to bind RBD with the mutations that appear in the current SARS-CoV-2 VOCs, B.1.1.7 (N501Y) and B.1.351 (K417N/E484K/N501Y), we screened the mutated RBDs by ELISA (Figure 3A, B). None of the mutations completely prevented recognition by the antibodies. Individual mutations N501Y, K417N and E484K, and N439K mutation (widespread in Europe), had only a marginal impact on the antibody binding. The combination K417N/E484K/N501Y exhibited either no or only minor influence on the binding of the antibodies of the Epitope I group. The mutations showed practically no influence on antibodies AX99, AX677 and AX97 of the Epitope II group. E484K, however, reduced binding of AX322, AX175 and partially also AX67 to RBD. E484K and N501Y made the highest impact on the binding of these mAbs to RBD containing the triple mutation N501Y/E848K/K417N present in the South African variant B.1.351.

**Figure 3.**
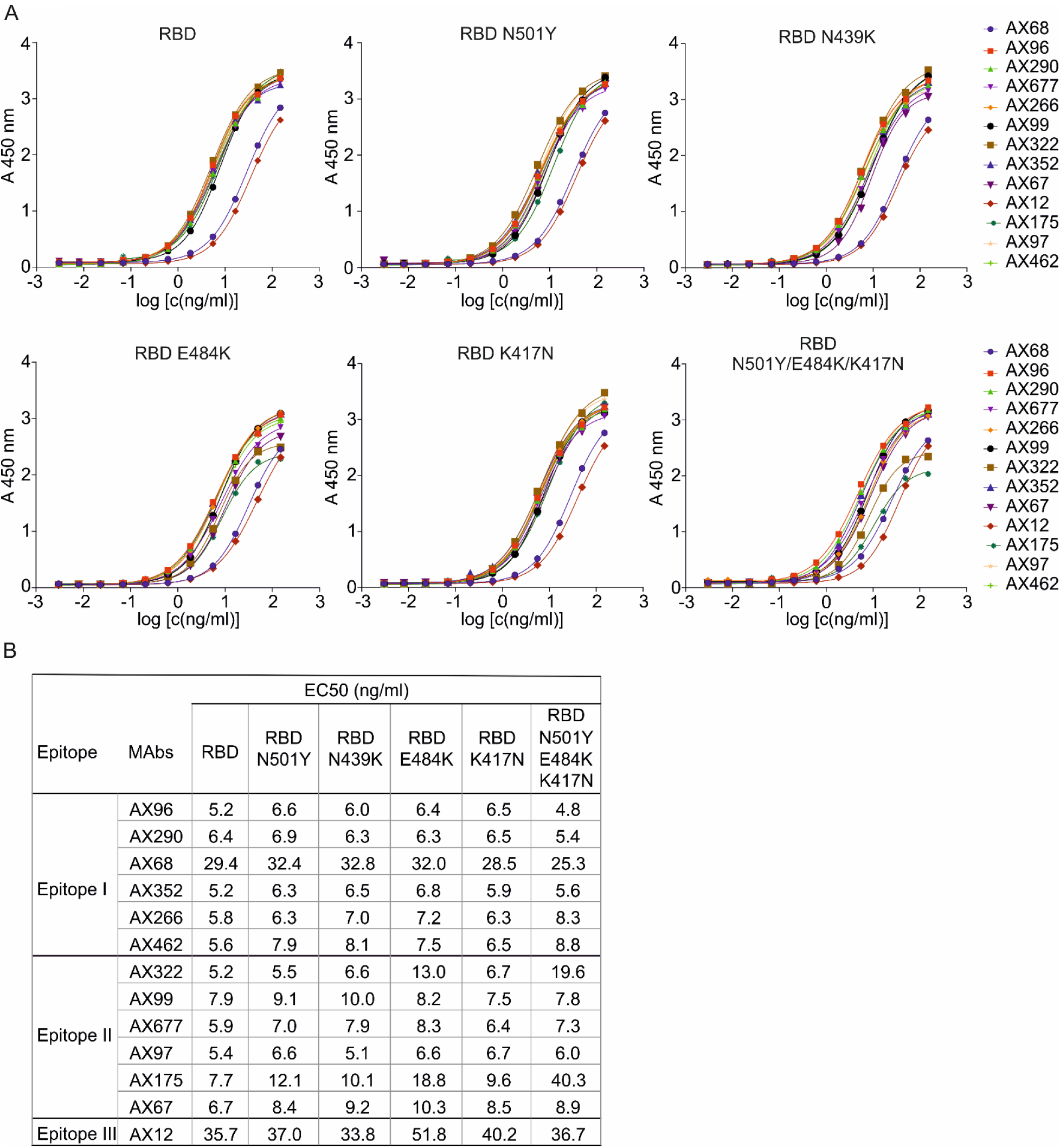
Effect of RBD mutations on immunoreactivity of monoclonal antibodies in ELISA and PRNT. (A, B) Binding of the antibodies to the RBD carrying the individual mutations (N501Y, N439K, E484K, K417N) and RBD with triple mutation N501Y/E484K/K417N were analysed. Effectivity of binding for each antibody was expressed by EC50 values.

Selected mAbs resistant against RBD mutations were tested for their neutralization activities in the plaque reduction neutralization test using the live authentic SARS-CoV-2 virus B.1 and its VOCs B.1.1.7 and B.1.351 (Figure 4). AX96 and AX290 effectively neutralized all three virus variants. AX677 was the third best neutralizing antibody, highly efficient against the wild type and B.1.351 viruses, with slightly reduced activity against B.1.1.7. Interestingly, AX68 showed relatively high activity against the B.1.351 variant with triple mutation in RBD, but 5- to 7-fold lower activity against the wild type and B.1.1.7, respectively.

**Figure 4.**
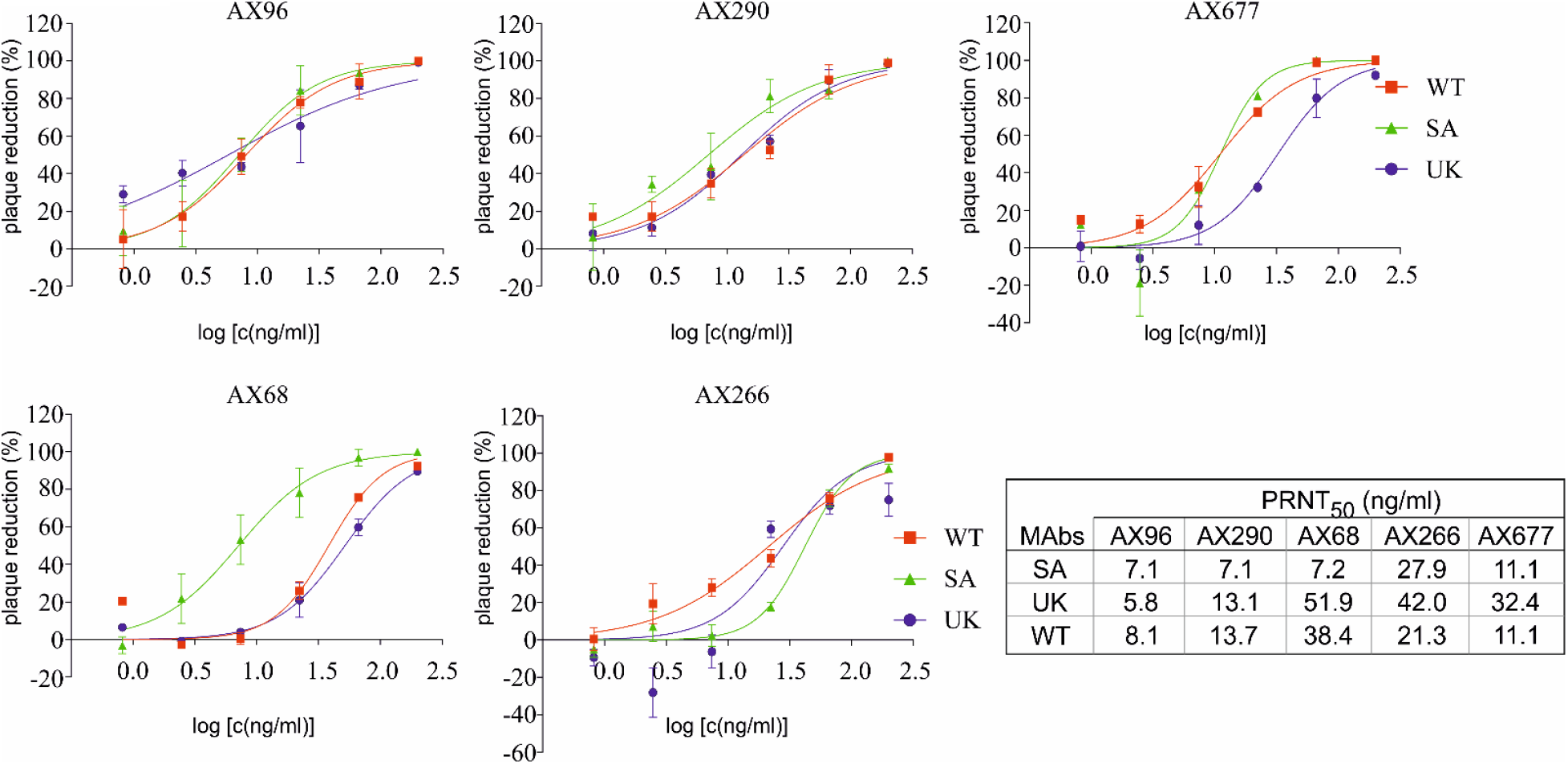
Selected mouse monoclonal antibodies neutralize SARS-CoV-2 variants of concern. Live authentic SARS-CoV-2 virus variants: Slovakia/SK-BMC5/2020 isolate (WT), B.1.1.7 (UK) and B.1.351 (SA), were pre-incubated with serial dilutions of Abs (250 ng/ml – 2 ng/ml) and then added to VERO E6 cells. Plaque reductions (%) relative to negative control were calculated 72 h post infection. Data are plotted as the mean from two wells of one experiment. PRNT_50_ mAb concentrations are shown in the table.

### Combination of antibodies with non-overlapping epitopes prevents mutational escape of the authentic SARS-CoV-2 virus

SARS-CoV-2 virus has accumulated mutations resulting in compromised activities of some vaccines and therapeutic antibodies ^30,31^. We, therefore, examined the ability of the authentic virus to escape from the selection pressure of AX290 and AX677, the best neutralizing mAbs with non-overlapping epitopes, alone and in combination (Figure 5A, B).

**Figure 5.**
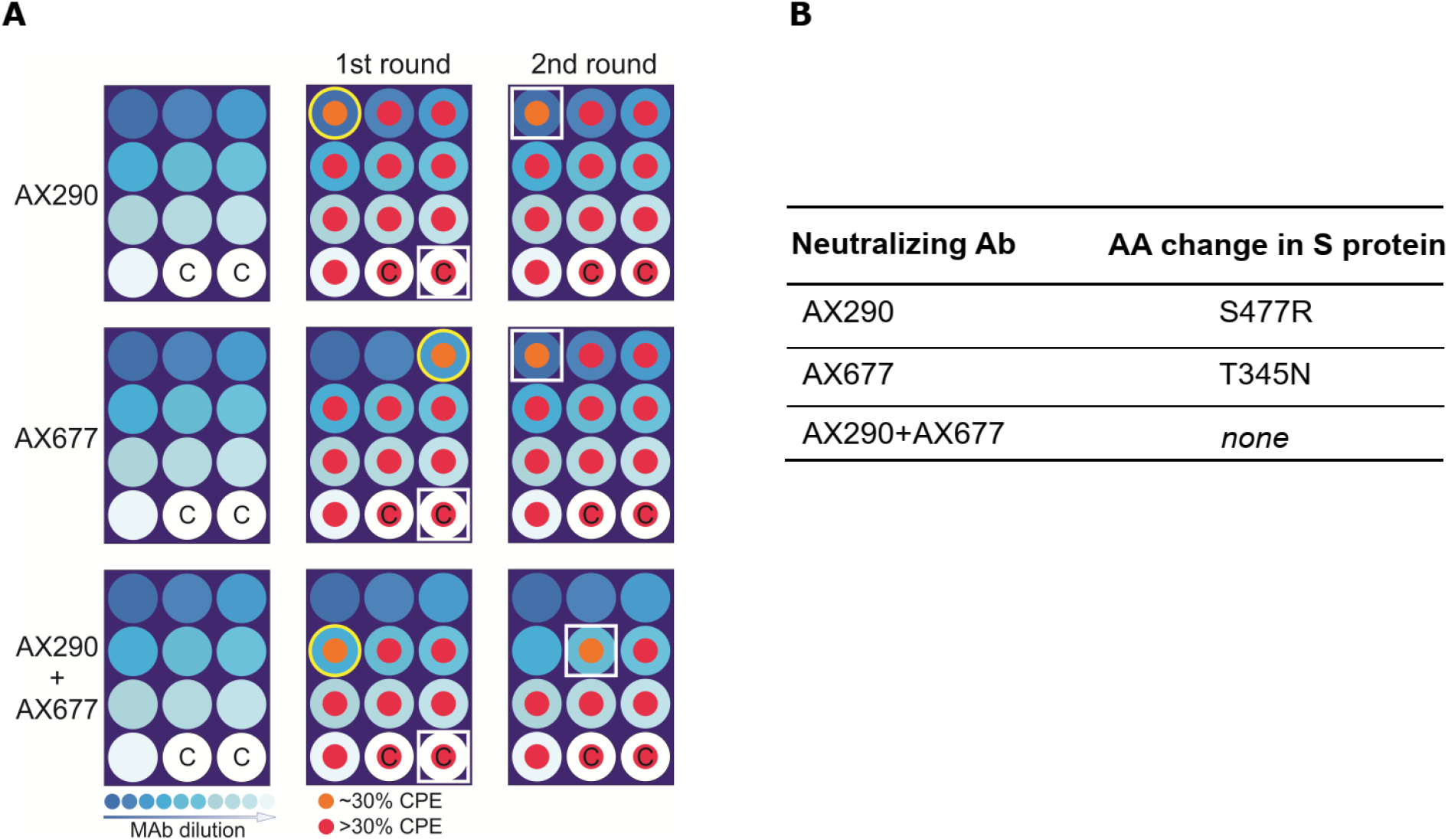
A combination of AX290 and AX677 prevents rapid mutational escape of authentic live SARS-CoV-2 virus. (A) A schematic diagram of microplate well allocations in the SARS-CoV-2 escape mutants experiment. MAbs were serially five-fold diluted starting with 50 μg/mL (left panels), pre-incubated with live SARS-CoV-2 (Slovakia/SK-BMC5/2020) virus (diluted to MOI=0.5) and added to VERO E6 cells (1^st^ passage, middle panels). Culture medium from the first wells of the mAb dilution series where CPE appeared (indicated with the yellow circles) were used for the 2^nd^ passage (right panels). White squares highlight wells with CPE where virus genomes were sequenced. C - control wells with virus without mAbs were sequenced to monitor possible tissue culture adaptations. (B) The table shows amino acid changes resulting from nonsynonymous mutations in the virus S protein that appeared under selection pressure of mAbs. Combination of the two antibodies prevented appearance of escape mutations.

It has been shown that an S-typed pseudoviruses might not fully recapitulate the complex behaviour of the authentic SARS-CoV-2 as measured by neutralizing activities of tested antibodies ^32^. We have also observed that our highly neutralizing antibodies behave differently on S-pseudotyped mouse leukemia virus (MLV), AX677 did not neutralize it while AX290 did (Figure 1C). Therefore, we have used live authentic virus isolate Slovakia/SK-BMC5/2020, that represents strains circulating in Europe in spring 2020, for the escape test. The virus particles, preincubated with serially diluted antibodies (individual or their 1:1 mixture), were used to infect Vero E6 cells. When the cytopathic effect of the escaping viruses became evident, the cell culture media with the escaping viruses (wells with approx. 30% cells affected) were collected, again preincubated with the respective serially diluted antibodies and added to fresh Vero E6 cells for another round of mutant virus amplification (Figure 5A). Sequence analysis of the viruses that escaped from the neutralization by the individual antibodies revealed only one mutation in the Spike protein per antibody: a S477R substitution for AX290, and a T345N substitution for AX677 (Figure 5B).

The S477R substitution that allowed escape of the virus from AX290 is located in the region of RBD that was identified by HDX as the contact site of the antibody (Figure 5C). It was found in 0.0705% of the sequenced SARS-CoV-2 isolates so far (https://www.gisaid.org/hcov19-mutation-dashboard/, database version as of May 23, 2021, 1621641 sequences analysed). The alternative substitution S477N that is more widespread in circulating viruses (3.31%) was not found in the sequenced viral genomes.

The mutation T345N present in the escape virus mutant against AX677 is positioned close to peptides protected by the antibody, but not directly within their sequence. It is a very rare mutation, only one T-N substitution at the position 345 of Spike has been found in circulating viruses, and only 32 substitutions of T345 were detected in the GISAID database until now. It is possible that this region of Spike RBD is less targeted by human immune repertoire resulting in a low selection pressure, or mutations there compromise the viral fitness. The T345N escape mutation has also been identified previously using an S-typed VSV pseudovirus for antibody 2H04 ^33^, which also did not compete with ACE2 for binding to the S protein.

Importantly, no escape mutation in the Spike protein was detected with the antibody mixture. The cytopathic effect appeared simply due to the dilution of the antibody concentration below the threshold required to neutralize the virus (625-fold higher dilution than any single mAb in the second passage). This shows that the combined therapy with both AX290 and AX677 would provide a potent barrier impeding viral replication and would protect against the mutational escape.

### Mouse-human chimeric versions of AX290 and AX677 maintain their potent neutralizing activities against authentic SARS-CoV-2 variants of concern

In order to prepare AX290 and AX677 for clinical testing, their variable regions were fused to human kappa and IgG1 constant regions. The chimeric mAbs AX290ch and AX677ch exhibited similar binding and functional properties as parental mouse versions. Both chimeric antibodies bound various mutant forms of RBD in ELISA with similar activities (Figure 6A).

**Figure 6.**
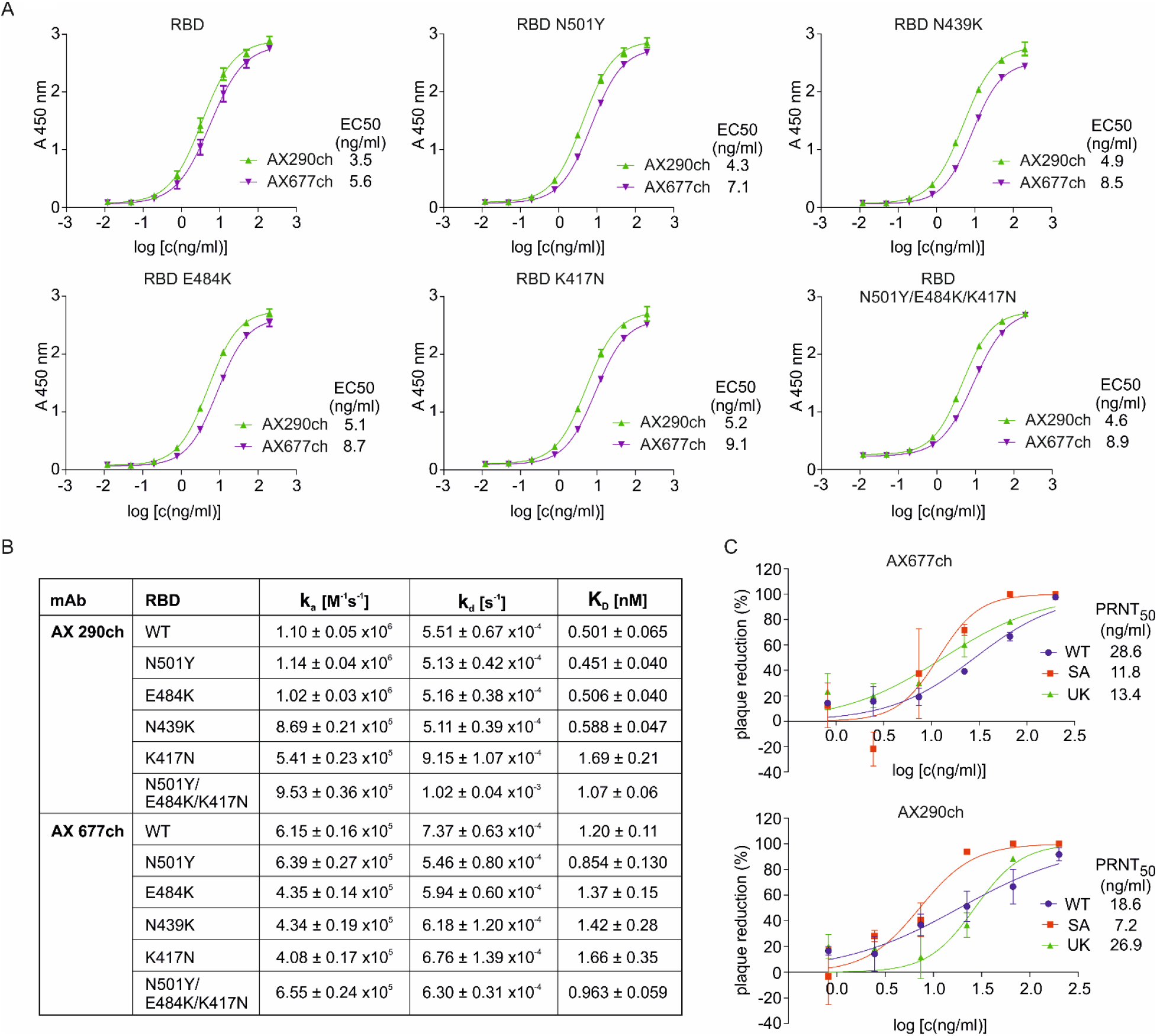
Functional characteristics of chimeric monoclonal antibodies. (A) Binding of the chimeric antibodies AX290ch and AX677ch to the Spike protein RBD and RBD carrying the individual and triple mutations were analysed by ELISA and binding to each antibody was expressed by EC50 value. (B) Neutralization activity of the two chimeric antibodies against live authentic SARS-CoV-2 virus variants: Slovakia/SK-BMC5/2020 isolate (WT), B.1.1.7 (UK) and B.1.351 (South Africa), determined in plaque reduction neutralization test (%). Data are plotted as the mean from two replicates within one experiment. (C) Kinetic characteristics of the interactions of WT and mutant RBD with selected neutralizing chimeric antibodies.

The chimeric antibodies recognised all tested mutants with nanomolar or sub-nanomolar affinities, which guarantees their unmodified neutralizing capacity (Figure 6B). Numerically, AX290ch maintained its picomolar affinity to all mutants except those containing K417N mutation alone or in combination N501Y/E484K/K417N (3-fold and 2-fold decrease in the affinity, respectively). AX677ch binds N501Y mutation, with 1.4-fold higher affinity than wild type RBD, reaching picomolar values (Figure 6B). Furthermore, this positive effect of the N501Y mutation is propagated into the triple mutant combining mutations N501, E484K, and K417N, which AX677ch binds with higher affinity than either of the two single mutants E484K and K417N, and even wild type RBD (1.4-, 1.7- and 1.2-fold, respectively).

The chimeras efficiently neutralized live authentic SARS-CoV-2 virus, Slovakia/SK-BMC5/2020 isolate (B.1 lineage), and variants of concern B.1.1.7 and B.1.351, in PRNT (Figure 6C). The tests confirmed that the neutralizing activity was fully preserved in the chimeric versions of the antibodies.

## Discussion

Monoclonal antibodies targeting the viral protein S have enormous potential to prevent SARS-CoV-2 infection and treat patients with mild to moderate COVID-19 ^34^. Several antibodies targeting RBD of the Spike protein have already been authorised for emergency use, including REGN10933/REGN10987, LY-CoV555 and JS016/LyCoV016 ^1^ and VIR-7831 with others in the pipeline (TY027, CT-P59, BRII-196 and BRII-198, SCTA01 etc.). The emergence of new SARS-CoV-2 variants of concern B.1.1.7, B.1.351, and P.1, with mutations in Spike protein and increased human-human transmissibility ^20–23^, raised concern about the efficacy of vaccines and therapeutic antibodies ^26,30,31^.

It was indeed shown that SARS-CoV-2 variants B.1.351 and P.1 were partially (REGN10933) or completely (LY-CoV555) resistant against antibodies used for COVID-19 treatment ^30^. This is in concordance with a study showing that the interaction between the mutant RBD containing three mutations N417/K484/Y501 and LY-CoV555/Bamlanivimab was completely abolished ^31^. To avoid this therapeutic limitation, we screened and selected the antibodies showing neutralization activity against several variants of SARS-CoV-2.

We took advantage of the mouse immunoglobulin repertoire ^18^ through the hybridoma technology to isolate monoclonal antibodies neutralizing live authentic SARS-CoV-2 virus. The *in vitro* and live virus screening assays allowed us to identify two neutralizing antibodies, AX290 and AX677, each of them neutralizing live authentic SARS-CoV-2 virus circulating in Europe in the spring of 2020 and two fast-spreading authentic variants of concern B.1.1.7 and B.1.351. It is important to emphasize that the antibodies were resistant to the E484K mutation, which has been recently shown to endow the S-typed VSV pseudovirus with resistance to several human convalescent sera ^33^. The authors have suggested that the repertoire of antigenic sites on RBD is limited in some individuals and that this amino acid is a part of a dominant neutralizing epitope.

The use of a live authentic SARS-CoV-2 virus to identify the escape mutations, as opposed to other studies that employed an S-typed pseudovirus ^10,33,35–38^, allowed to eliminate a potential bias from an inadequate infection and replication machinery of a model virus. It has been noted that only some of the spike escape mutations identified in SARS-CoV-2 S-typed infectious vesicular stomatitis virus were found so far in the context of authentic SARS-CoV-2 virus isolates, it is possible that those not identified might compromise the fitness of the genuine virus^33^.

Our authentic virus approach identified antibody-specific escape mutations in RBD of the Spike protein, S477R and T345N, that directly affect the recognition by the antibodies.

The combination of the two antibodies exhibits a strong synergistic neutralizing effect against the authentic live SARS-CoV-2 virus observable at >600-fold lower concentration than each antibody alone. More importantly, the antibody combination also prevented emergence of escape mutations of the authentic SARS-CoV-2 virus.

The potential for an effective COVID-19 treatment and protection against antibody resistance is strengthened by the fact that they bind non-overlapping epitopes on functionally different portions of the S protein RBD and apparently different mechanisms of how the two antibodies neutralize the virus. In addition, the chimeric antibodies AX290ch and AX677ch showed similar binding potencies against various mutations located in RBD and effectively neutralized SARS-CoV-2 variants of concern.

### Limitations of the study

We recognize that the resistance of the antibody cocktail against the virus escape mutations was not confirmed in PRNT. The mechanism of virus neutralization by AX677 has not yet been understood.

## Acknowledgements

The study was funded by AXON Neuroscience SE.

## Author contributions

EK, MZ, NPI, LF, NTC, MP, DP, KM performed primary screening of antibodies and in vitro assays. RS, OC, KT performed affinity measurement. PF, MC, MS, GPR, KS, NB, JP generated and produced recombinant proteins and chimeric antibodies. BKl, MS performed plaque reduction neutralization assay and virus escape. KB, VC, BB, TV, JN performed sequence analysis of the viruses and bioinformatics. AK, PM, VP, JP performed proteomic analysis. BKo, NZ, MN, MF, EK conceptualized and designed experiments, supervised the study and prepared the manuscript.

## Materials and Methods

### Cell lines

Human embryonic kidney HEK293T/17-hACE2 cells with stable expression of human angiotensin-converting enzyme ACE2 (AXON Neuroscience SE) were prepared by stable transfection of pDUO2-hACE2-TMPRSS2a (InvivoGen) with transfection reagent Lipofectamine 3000 (Thermo Fisher Scientific), according the to the manufacturer’s recommendations. The hygromycin resistant colonies of stable transfectants were isolated and screened for the expression of the ACE2 protein by Western blotting. The expression of the serine protease TMPRSS2 was not evaluated. Stable cell line HEK293T/17-hACE2 were maintained in selection medium: Dulbecco’s Modified Eagle Medium (DMEM) supplemented with 10% heat-inactivated fetal bovine serum (FBS), 2 mM L-glutamine (GIBCO), 5μg/ml gentamicin (SIGMA) and 100μg/ml hygromycin B (Invitrogen) at 37°C in 5%CO_2_. In PRNT, Vero E6 cells (Vero C1008, ATCC CRL 1586) were cultured in Eagle’s minimal essential medium (EMEM, Lonza) supplemented with 5% FBS (GIBCO), Penicillin-Streptomycin-Amphotericin B Solution (10ml/l, Lonza, Switzerland).

### Proteins

The prefusion-stabilized SARS-CoV-2 S protein ectodomain (residues 1−1208 from GenBank: MN908947, with proline substitutions at residues 986 and 987, a “GSAS” substitution at the furin cleavage site residues 682–685, a C-terminal T4 fibritin trimerization motif, an HRV3C protease cleavage site, a TwinStrepTag and an 8XHisTag), the receptor binding domain (RBD) fragment of the S protein (amino acids 319-591), containing the His-tag and Twin-Strep-tag and modified ACE2 (angiotensin-converting enzyme 2), containing tags for purification were synthesized at BioCat Co. (BioCat GmbH, Germany).

### Viruses

Three cell culture isolates of SARS-CoV-2 were used within the study. They were all isolated from clinical samples of COVID-19 patients in Slovakia. All three isolates were deposited in the European virus archive GLOBAL. The strain Slovakia/SK-BMC5/2020 (available at https://www.european-virus-archive.com/virus/sars-cov-2-strain-slovakiask-bmc52020) represents strains circulating in Europe in spring 2020 and carries the Spike D614G mutation (lineage B.1). The strain Slovakia/SK-BMC-P1A/2021 (available at https://www.european-virus-archive.com/virus/sars-cov-2-strain-slovakiask-bmc-p1a2021-b117-variant-voc-20201201) was isolated in January 2021 and belongs to the B.1.1.7 lineage (VOC 202012/01). The strain Slovakia/SK-BMC-BA11/2021 (available at https://www.european-virus-archive.com/virus/sars-cov-2-strain-slovakiask-bmc-ba112021-b1351-variant-aka-20h501yv2-or-south-african-variant) was isolated in March 2021 and belongs to the B.1.351 lineage (20H/501Y.V2). The complete genome sequences of all three viruses were deposited in the GISAID.org database under the accession IDs EPI_ISL_417879, EPI_ISL_77965.1, and EPI_ISL_ 1234458, respectively.

### Preparation of plasmids coding for S, S-RBD and ACE2 proteins

For protein production, the synthetic genes (BioCat) coding for the Spike protein (S), receptor binding domain (RBD) or ACE2 were cloned under the CMV promoter in a pCMV-based mammalian expression vector immediately after a signal peptide accomplishing secretion of recombinant proteins from the cells into medium. The plasmids were amplified in DH5α bacterial cells. The cells carrying the plasmid were inoculated into 5 ml LB medium with kanamycin for 8 hrs, transferred into 250 ml cultures and incubated overnight with shaking at 37°C. The plasmid DNA was isolated using PureLink® HiPure Plasmid Filter Maxiprep Kit (Invitrogen) with the last ethanol washes done in a sterile laminar-flow hood. After overnight incubation in sterile water at 4°C, DNA concentration was measured using NanoPhotometer and the DNA was aliquoted and stored at −20°C. Insertion of the desired point mutations in S-RBD was done using QuickChange II site directed mutagenesis kit (Agilent Technologies). Primers for mutagenesis were designed using The QuickChange® Primer Design Program provided by manufacturer. Prepared mutations, verified by DNA sequencing were transformed in DH5α bacterial cells for further amplifications.

### Production of the proteins in human cell line Expi293

Proteins were produced in human cell line Expi293. Plasmids that drive protein expression from the CMV promoter were either transfected with polyethyleneimine (PEI) or by electroporation into Expi293 cells that were in exponential growth phase. For PEI transfections, 3.75ml PEI and 1.25mg DNA was added to a suspension of 10^9^ cells in a total volume of 50 ml, shaken at 37°C for 3 hrs and diluted into working concentrations. For electroporation, 100 μg/ml DNA was electroporated in a suspension of 2.10^8^ cells/ml in electroporation buffer using MaxCyte STX (MaxCyte, MD, USA), the cells were let to rest for 30 min and then diluted into the final concentration of 3.10^6^ cells/ml. The cultures were incubated with shaking at 37°C for 24 hrs, transferred to 32°C and further cultured until cell viability decreased to <50% (approx. 4-6 days), at which point the medium was harvested by centrifugation at 300*xg* for 12 min and stored at −20°C until purification.

### Protein purification

The proteins were purified from cell media using Åkta FPLC (GE Healthcare) as follows. The culture medium was clarified by centrifugation at 20,000*xg*, NaCl was added to a final concentration of 0.5 M and the solution was filtered through a 0.2 μm membrane filter. A 5 ml His-Trap affinity column (Cytiva) charged with nickel ions was equilibrated in a 20 mM sodium phosphate buffer, pH 7.4, 0.5 M NaCl and the culture medium was loaded onto the column. Subsequently, the column was washed with 20 ml of the 20 mM sodium phosphate buffer, pH 7.4, and His-tagged protein was eluted by 0.5 M imidazole in 20 mM sodium phosphate buffer, pH 7.4. The fractions containing the target protein were pooled and further purified on the Strep-Tactin® media (IBA GmbH, Germany) as described by the manufacturer, using 10 mM desthiobiotin as an elution reagent. Purified proteins were concentrated by ultrafiltration and buffer-exchanged into PBS on a 5 ml HiTrap Desalting column (GE Healthcare). The concentration was determined from UV absorbance at 280 nm. The proteins were sterile-filtered and stored at −20°C. Extracellular part of the human ACE2 protein fused to human IgG Fc fragment, preceded by a HRV3C protease cleavage site was purified by His-Trap and Strep-Tactin® media as above. The Fc fragment was cleaved out by an overnight incubation with HRV3C protease at +6°C. ACE2 was afterwards polished by size-exclusion chromatography on a Superdex 75 16/60 column (GE Healthcare), concentrated by ultrafiltration with a Amicon Ultra centrifugal filter of 10 kDa MW cut off (Millipore), sterile-filtered and stored at +6°C.

### Preparation of hybridoma cell lines producing monoclonal antibodies specific to SARS-CoV-2 spike protein and RBD

Six-week-old Balb/cN mice were primed subcutaneously either with 30 μg of recombinant SARS-Cov-2 Spike protein (S) or Spike-RBD (RBD) in the complete Freund’s adjuvant (SIGMA-ALDRICH) and boosted three times at three-week intervals with 20 μg of the same antigen in the incomplete Freund`s adjuvant. Three days before the spleen extraction, mice were injected intraperitoneally with 20 μg of the same antigens in PBS. Spleen cells from immunized mice were fused with NS0 myeloma cells according to the method of Kontsekova *et al.^39^*. Splenocytes were mixed with NS0 myeloma cells (ratio 5:1) and fused for 1 minute in 1 ml of 50% polyethylene glycol (PEG) 1550 (Serva) in serum free Dulbecco’s modified Eagle’s medium (DMEM) supplemented with 10% dimethyl sulfoxide (DMSO). The fused cells were resuspended in DMEM containing 20% horse serum, L-glutamine (2 mM), HAT media supplement Hybri-Max™ (SIGMA), and gentamycin (40 U/ml), at a density of 2.5 × 10^5^ spleen cells per well in 96-well plates and incubated for 10 days at 37°C.

### ELISA for screening of monoclonal antibodies

The growing hybridomas were screened for the production of monoclonal antibodies specific to the S protein and RBD by an enzyme-linked immunosorbent assay (ELISA). Microtiter plates were coated overnight with one of the proteins (2 μg/ml, 50 μl/well) at 37°C in PBS. After blocking with PBS-0.1% Tween 20 to reduce nonspecific binding, the plates were washed with PBS-0.05% Tween 20 and incubated with 50 μl/well of hybridoma culture supernatant for 1 hr at 37°C. Bound monoclonal antibodies were detected with goat anti-mouse immunoglobulin (Ig) conjugated with horse radish peroxidase (HRP, Dako, Denmark). The reaction was developed with TMB one (Kementec Solutions A/S, Demark) as a peroxidase substrate and stopped with 50 μl of 0.25 M H2SO4. Absorbance at 450 nm was measured using a Powerwave HT (Bio-Tek). Readouts with an absorbance value of at least twice the value of the negative controls (PBS) were considered positive. ELISA positive hybridoma cultures were further subcloned in soft agar according to the procedure described previously^39^. After purified hybridoma clones were obtained, the antibodies isotypes were determined by ELISA using a mouse Ig isotyping kit (ISO-2, SIGMA).

### Determination of kinetic characteristics of the antibody-RBD interaction by surface plasmon resonance (SPR)

Kinetic characteristics and affinities of mouse and chimeric antibodies to RBD was measured by SPR. A BIACORE3000 instrument with a CM5 sensor chip (Cytiva, USA) was used for the SPR assays. Amine-coupling reagents (EDC, NHS, ethanolamine pH 8.5), were obtained from Cytiva. Typically, 10 000 RU (response units) of polyclonal anti-mouse antibody (No. Z 0420; Dako, Denmark) or polyclonal anti-human antibody (I2136, Merck, Germany) was coupled simultaneously in four flow cells. One of them was used as a reference cell in kinetic measurements. The experiments were done at 25°C in PBS (pH 7.4) supplemented with 0.005% of Tween 20 (PBS-P). In each analysis cycle, three different antibodies were captured in three analytical flow cells to reach an immobilization level of 100-150 RU. Subsequently, four serial dilutions of RBD protein ranging from 3.125-25 nM or the running buffer (PBS-P) as a control were injected in duplicates at a flow rate of 100 μl/min over the sensor chip. Regeneration of the chip surface was accomplished by a 6 s injection of 100 mM HCl. Kinetic binding data were double referenced by subtracting the reference cell signal and the PBS-P buffer response and fitted by BIA evaluation software 4.1 (Biacore AB) to a 1:1 interaction model as described previously^40^. On- and off-rates were fitted locally, maximal response was fitted globally and the bulk response was set to zero. The equilibrium dissociation constants KD of antibody-RBD complexes were calculated from the averaged association and dissociation rate constants. The final error of affinities was propagated from errors of the experimental rate constants used for KD calculation.

### Epitope binning by competitive ELISA

For analysis of cross-competition between RBD/S specific mAbs, competitive ELISA was developed. Fab fragment of anti-mouse IgG (0.5 μg/ml, 50 μl/well in PBS) was immobilized overnight on microtiter ELISA plate at 37°C. After blocking with PBS-0.1% Tween 20 to reduce nonspecific binding, the plates were washed with PBS-0.05% Tween 20 and incubated with 50 μl/well of monoclonal antibody 1 (0.6 μg/ml) for 1hr at 37°C. Horseradish peroxidase (HRP)-conjugated RBD, at a concentration of 20 ng/ml, was pre-incubated with monoclonal antibody 2 (at concentrations of 0.25 – 2 μg/ml) for 1 h at room temperature, then the mixture was added to the ELISA plate and incubated at 37°C for 1h. Binding of HRP-conjugated RBD to immobilized antibody 1 was detected with TMB one substrate (Kementec Solutions A/S, Demark) and stopped with 50 μl of 0.25 M H_2_SO_4_. Absorbance at 450 nm was measured using a Powerwave HT (Bio-Tek) plate reader.

### RBD-ACE2 binding inhibition assay

With the aim to select the antibodies blocking the interaction of receptor binding domain (RBD) with angiotensin-converting enzyme 2 (ACE2) receptor, ELISA-based inhibition test was used. 96-well microtitre plates (Nunc MaxiSORP) were coated with ACE2 recombinant protein at 350 ng per well in 50 μl of phosphate-buffered saline (PBS) overnight at 4°C, followed by blocking with PBS supplemented with 0.1% Tween-20 (PBS-T). Horseradish peroxidase (HRP)-conjugated RBD at a concentration of 0.4 μg/ml was pre-incubated with tested hybridoma culture supernatant producing mAbs (diluted 1:6 with PBS-T) for 1 h at room temperature (final volume of 50 μl), followed by addition into the ELISA plates coated with ACE2 and incubated for 1 h at 37 °C. Unbound HRP-conjugated RBD were removed by five washes with PBS-T. A colorimetric signal was developed by the enzymatic reaction of HRP with a chromogenic substrate TMB one (Kementec Solutions A/S, Denmark). An equal volume of stop solution (0.25 M H_2_SO_4_) was added to stop the reaction, and the absorbance of the product at 450 nm was measured using a PowerWave HT microplate reader (BioTek). Inhibition activity of antibody was determined as follows:

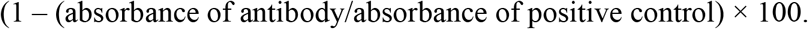

### S-ACE2 binding inhibition assay

HEK 293T/17 cells stably expressing human ACE2 protein (HEK 293T/17-hACE2) were seeded at 60-70% plating density in 48-well plates and cultivated O/N at 37°C, 5% CO_2_ in a humidified incubator in DMEM supplemented with 10% (v/v) fetal calf serum, 2 mM L-glutamine, 100 units/mL penicillin/streptomycin (all from Life Technologies Invitrogen, Carlsbad, CA, USA) and 100 μg/ml hygromycin (Thermo Fisher Scientific). Recombinant Spike protein was labelled with Alexa Fluor™ 546 (Thermo Fisher Scientific) according to the manufacturer’s recommendations. Labelled S protein (40 ng/ml) was preincubated with tested hybridoma culture supernatants producing mAbs (diluted 1:50 in DMEM) for 30min at 37°C. Then the preincubation mixtures were added to HEK 293T/17-hACE2 cells and incubated for 2 hrs at 37°C, 5% CO_2_ in a humidified incubator. Subsequently, cells were gently re-suspended in 500 μl PBS, transferred into flow cytometry tubes and immediately evaluated for S protein internalization by flow cytometry. Cells were gated using forward and side scatter to exclude cellular debris. The FSC-H/FSC-A gating approach was used to eliminate doublets.

Measurements were recorded as mean fluorescent intensity of Alexa Fluor 546. Approx. 10 000 cells were quantified for each microtiter plate well. Fluorescence data were normalized to cells treated with the labelled S protein pre-incubated with a hybridoma culture supernatant from a clone producing an irrelevant antibody. IC50 values for particular experiments were calculated by nonlinear regression analysis using GraphPad Prism (GraphPad Software Inc.).

### Pseudovirus SARS-CoV-2 neutralization assay

For testing the neutralization ability of monoclonal antibodies, the procedure adapted according to Nie et al., (2020) was used. Briefly, the recipient cells HEK293/17-hACE2 were seeded in 96-well plate and cultured overnight in a humidified CO_2_ incubator (at 5% CO_2_ and 37°C) in 100 μL of normal growth medium (DMEM) with 10% of FCS. In parallel, the optimal S-typed pseudoviral particles (S-PVPs) dilution was determined. “SARS-CoV-2 Pseudoviral Particles Spike” from MyBiosource (San Diego, CA, USA) were used in the assay (Cat. No. MBS434275). PVPs concentration of the material was ≥ 1×10^7^ (as determine from TEM scan by the manufacturer). PVPs were aliquoted and stored at −80°C.

Optimal dilution of the original PVPs stock for testing of the antibody neutralization efficiency was set in the range from 1:100 to 1:200, since this dose results in 50% of recipient cells infected with S-PVPs. The neutralization assay was performed as follows: the serially diluted selected monoclonal antibodies were mixed with a defined amount of S-PVPs. The mix was incubated at 37°C in a CO_2_ incubator for 1 hr. Then the mix was added to the cells in a 96 well–plate (50 μl of mix per well) in triplicates. After 48 hrs of incubation of the recipient cells with the S-PVP mix the luciferase activity was measured using a luminometer (Fluoroskan Ascent® FL, Labsystems) and half maximal effective concentration was calculated for the measured monoclonal antibodies. The measurement of relative luminescent units (RLU) and comparison to the positive control (antibody not recognizing S protein) and a negative control (cell extracts with no S-PVPs or extracts from the cells infected with S-deficient PVPs) was performed in all experiments.

### SARS-CoV-2 plaque reduction neutralization assay

The plaque reduction neutralization test (PRNT) with the live SARS-CoV-2 virus (performed in a containment laboratory of Biosafety level 3 by the Department of Virus Ecology at the Biomedical Research Center of the Slovak Academy of Sciences, Slovak Republic) was used as a test for determination of neutralization capacity for the coronavirus of mAbs elicited by immunization with the Spike protein or its RBD. Serial dilutions of supernatants from the hybridoma clones producing the selected mAbs were incubated with 100 plaque-forming units of SARS-CoV-2 at 37°C for 2 hrs. MAb–virus mixtures were then added to Vero E6 cell monolayer in 24-well plates and incubated at 37°C for additional 1 h. After incubation, cells were overlaid with 2% (w/v) carboxymethylcellulose in Eagle’s minimal essential medium (EMEM) supplemented with 5% FBS. Plates were incubated at 37°C for 72 hrs. Then, the cells were fixed for 30 min with 4% formaldehyde in phosphate-buffered saline (PBS). After incubation the SARS-CoV-2 plaques were visualized by staining with 0.5% crystal violet at room temperature for 10 min. After washing the wells with water, the number of plaques was counted. The antibody neutralization activity was determined as the reciprocal of the highest dilution resulting in an infection reduction of 50% (PRNT_50_). Data were fitted to logistic 4-parameter sigmoidal dose response curve using GraphPad Prism (GraphPad Software Inc.), the goodness of fit was R^2^>0.9, except for AX266 on B1.1.7 (R^2^=0.7888), AX96 on B.1.1.7 (R^2^=0.8766), AX290ch on B1.1.7 (R^2^=0.8730) and AX677ch on B.1.351 (R^2^=0.8598).

### Virus escape from neutralizing antibodies

For the first round of the escape mutant selection, SARS-CoV-2 virus at a high multiplicity of infection (MOI) of 0.5 was added to each dilution of mAbs (5-fold serial dilution starting at 50 μg/mL) and incubated at 37°C for 1hr. The mixture was then transferred to 12-well plates containing sub-confluent Vero E6 cell monolayer. Plates were incubated 3-4 days at 37°C and 5% CO_2_ and examined for the presence of a cytopathic effect (CPE). Supernatants from the first wells in the mAb dilution series with evident CPE (approx. 30% cells affected) were collected. For the second passage, these supernatants were diluted 1:5 in DMEM and pre-incubated for 1hr at 37°C in the presence of serially 5-fold diluted mAbs starting at 100 μg/mL. After incubation at 37°C for 1hr, the mixture was transferred to 12-well plates with Vero E6 cells and further incubated 3-4 days at 37°C. Total RNA, including the viral RNA, was extracted from the cells in the first wells of the mAb dilution series that show CPE and used for the sequencing.

### Nanopore sequencing of viral RNA

The sequencing libraries were constructed essentially as described in the Eco PCR tiling of SARS-CoV-2 virus with native barcoding protocol (Oxford Nanopore Technologies), except the amplicons spanning the SARS-CoV-2 genome sequence were generated using the ~2.5-kbp primer panel^41^ in which the rightmost primer pair was replaced by corresponding pair from the ~2.0-kbp panel^42^. The primers were custom synthesized by Microsynth. Briefly, purified RNA (8 μl) was converted into cDNA using a LunaScript RT SuperMix Kit (New England Biolabs) and used as a template in two separate amplification reactions generating odd- and even-numbered tiled amplicons. The PCR was performed using a Q5 Hot Start High-Fidelity 2X Master Mix (New England Biolabs) and the cycling conditions were: 30 sec at 98°C (initial denaturation), followed by 30 cycles of 15 sec at 98°C (denaturation) and 5 min at 65°C (combined annealing and polymerization). The overlapping amplicons were pooled and purified using 0.5 volume of AMPure XP magnetic beads (Beckman Coulter). The ends of amplicons were treated with a NEBNext Ultra II End repair / dA-tailing Module (New England Biolabs) and barcoded using an Native Barcoding Expansion 96 kit (Oxford Nanopore Technologies) and a Blunt/TA Ligation Master Mix (New England Biolabs). Barcoded samples (96) were pooled and purified using 0.5 volume of AMPure XP magnetic beads. The AMII sequencing adapter (Oxford Nanopore Technologies) was ligated to about 300 ng of barcoded pools using Quick T4 DNA ligase (New England Biolabs) and the library was purified using 0.5 volume of AMPure XP magnetic beads. About 90 ng of the library was loaded on an R9.4.1 flow cell and the sequencing was performed using a MinION Mk-1b device (Oxford Nanopore Technologies).

### Variant calling

Raw nanopore data was base called and demultiplexed using Guppy v.4.4.1. Barcodes at both ends were required in demultiplexing. Variants were called using the ARTIC pipeline (https://artic.network/ncov-2019/ncov2019-bioinformatics-sop.html), which internally filters reads based on quality and length, aligns them to the reference (Wuhan-Hu-1 isolate, NC_045512) using minimap2 (https://github.com/lh3/minimap2), trims primers, and calls variants using Nanopolish (https://github.com/jts/nanopolish). All variants reported in Table S1 were determined based on read coverage of at least 380. All new nonsynonymous mutations were supported by at least 90% of the analyzed reads overlapping a particular site. Note that due to higher error rates of nanopore sequencing, even fixed variants typically do not achieve 100% support.

### Immunocytochemistry

HEK293T/17-hACE2 cells were plated on cover glass pre-coated with rat-tail collagen, type I (Sigma-Aldrich, St. Louis, Missouri, United States) and cultivated for 24 h in DMEM with 10% FCS. Cells were fixed with 4% paraformaldehyde and labelled with anti-human ACE2 antibody at 10 μg/ml (cat. # 600-401-X59; Rockland Immunochemicals), followed by goat anti-rabbit IgG secondary antibody at 1/500 dilution (green; cat. # A-11008; Invitrogen). The samples were mounted in Fluoroshield™ medium with DAPI (Sigma-Aldrich). Images were captured by LSM 710 confocal microscope (Zeiss, Jena, Germany).

### Hydrogen Deuterium Exchange Mass Spectrometry (HDX-MS)

RBD and AX290 or AX677 were dissolved in non-deuterated PBS buffer (Phosphate Buffered Saline, pH 7.4) and digested individually with flow through an immobilized nepenthesin-2 column, 2.1 mm × 20 mm (Affipro, Praha, Czech Republic) for peptide mapping. Prior to the HDX experiments, RBD and the AX290 or AX677 were incubated for 1 h to allow complex formation. To initiate HDX, proteins were diluted with D_2_O buffer (10 mM phosphate buffer, D2O, pD 7.0). Different aliquots were submitted to HDX for three times: 0.3, 2, 20, and 120 min. The reaction mixture was then quenched by adding quenching buffer (6M urea, 2M thiourea and 0.4 M TCEP, pH 2.3). The deuterated mixture was then digested online by using the same nepenthesin-2 column and the resultant peptides were desalted with flow through on a trap column (Phenomenex UHPLC fully Porous Polar C18, 2.1mm, Torrance, CA, USA) with 0.4% formic acid in water. The eluted peptides were further separated on a analytical column (Luna® Omega 1.6 μm Polar C18 100 Å, 100 × 1.0 mm, Torrance, CA, USA), by using a gradient of acetonitrile and water with 0.1% formic acid. Peptides were introduced to a timsTOF (Bruker, Brno, Czech Republic) mass spectrometer for mass analysis. The peptide-level deuterium uptakes were calculated by using MS Tools software ^43^.

